# Avian influenza causes age-related mortality in a long-lived seabird

**DOI:** 10.1101/2025.10.30.685500

**Authors:** Wouter Courtens, Mónika Ballmann, Ruben Fijn, Morten Frederiksen, Susanne Kühn, Luc Lens, Mardik Leopold, Sander Lilipaly, Florian Packmor, Eric Stienen, Peter van Horssen, Ulrich Knief

## Abstract

Recently, highly pathogenic avian influenza (HPAI) A (H5N1) clade 2.3.4.4b viruses have caused mass mortality events in seabirds worldwide, raising concern for long-lived species with low reproductive rates. Using individual-level data from the 2022 mass mortality event in northwestern European Sandwich terns (*Thalasseus sandvicensis*), we show that older individuals were disproportionately more affected, while no sex bias was observed. This age-specific mortality likely removed the most experienced individuals from the population. Our findings highlight a previously underappreciated mechanism through which HPAI outbreaks may impair the resilience of long-lived avian populations.

## Main

Long-lived seabirds are characterized by delayed maturation, low reproductive output, and high adult survival^1^. Because of these life-history characteristics, factors that increase adult mortality have a greater negative impact on population size than reductions in reproductive success or juvenile survival^2,3,4^. When highly pathogenic avian influenza (HPAI) A (H5N1) clade 2.3.4.4b viruses became enzootic in northwestern Europe in 2021^5^, it caused massive die-offs of adult seabirds^6^. Globally, more than 100 species of seabirds – including several endangered taxa – have been affected by this panzootic^7^, with total mortality reaching into the millions of individuals^8^. While these numbers already raise concerns about the ability of many species to recover^9^, the long-term impacts and prospects for recovery will largely depend on which demographic components are most affected^10^.

Age- or sex-biased mortality can influence population dynamics in long-lived species, but the extent and direction of this effect depend on age-specific survival and reproduction patterns^11^. Older individuals often contribute disproportionately to population growth through greater breeding performance based on individual experience or selective survival of high-quality birds^12,13^. Additionally, sex-biased mortality can strongly affect populations, especially in seabirds with monogamous mating systems and obligate biparental care^14^. Understanding these demographic processes is essential for predicting population recovery after mortality events.

In 2022, the Sandwich tern (*Thalasseus sandvicensis*), a seabird with a recorded lifespan of more than 30 years, was one of the bird species severely affected by HPAI in northwestern Europe, losing over 20,500 adults (> 17% of its regional population^15^). The virus spread rapidly through the densely populated colonies, where intense contact between breeding as well as prospecting birds likely facilitated transmission, both within and between colonies^16^. Here, we test whether sex- or age-related mortality occurred in this species during the 2022 mass mortality event.

Sex-specific mortality was assessed by sexing 243 dead adult Sandwich terns of known age, collected in the Netherlands during the 2022 outbreak. We observed an almost equal sex ratio (123 females, 120 males), with no significant deviation from 1:1 (binomial test, *P* = 0.90).

We assessed age-related mortality by first estimating apparent survival rates before the 2022 mass mortality event (1995–2021, Figs. 1d and 1g). For this, we used age-structured Cormack-Jolly-Seber (CJS) capture-mark-recapture models on resightings of 50,646 individuals ringed in their first year of life (1Y) in two regions that were at the core of the HPAI outbreak: the southern Netherlands & Belgium and the northern Netherlands (Figs. 1c and 1f). This allowed us to predict the age distribution of ringed birds alive and present in these regions immediately before the outbreak in 2022 (Fig. 1e and 1h). During the outbreak, a total of 832 ringed birds of known age from the two study regions were recovered dead: 595 in Belgium and the Netherlands and 237 elsewhere (compared to a mean ± standard deviation of 5.30 ± 3.36 ring recoveries per year in the preceding 27 years). We quantified HPAI-related mortality in two ways: (i) the ‘regional mortality’ estimate includes only dead birds recovered in their natal region and is therefore conservative, analogous to the estimation of apparent survival in CJS models, (ii) the ‘total mortality’ estimate includes all European recoveries of Belgian and Dutch ringed birds, both in and outside their natal region, and thus provides a more comprehensive estimate of the total HPAI-related mortality; it likely remains an underestimate due to imperfect detection and reporting.

**Fig. 1.**
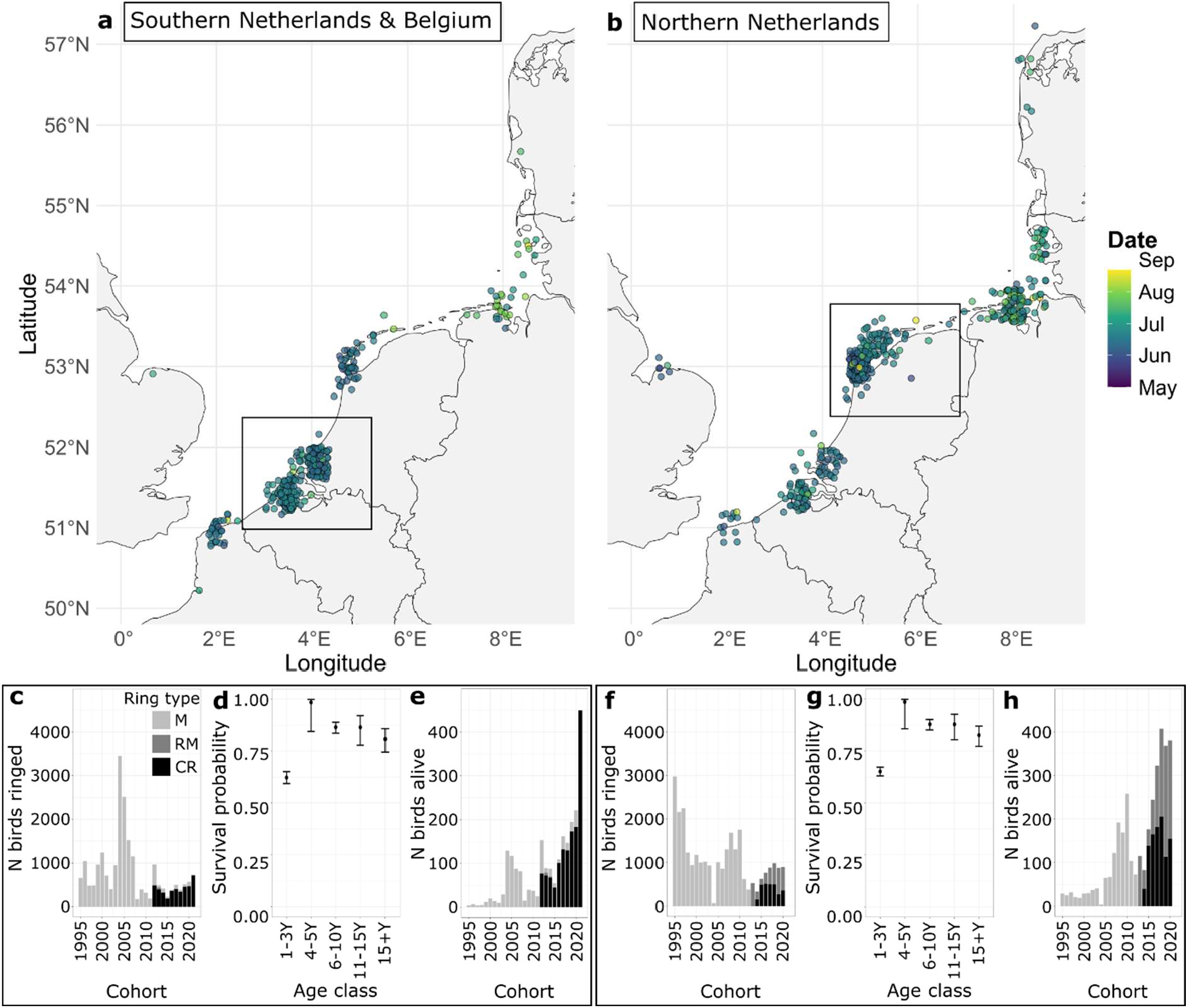
Distribution and demographic context of HPAI-related mortality in ringed Sandwich terns. **a & b**, Spatial distribution of dead Belgian/Dutch ringed Sandwich terns following HPAI A (H5N1) virus infection, May – August 2022, in the southern Netherlands & Belgium (a) and the northern Netherlands (b). Dots indicate individual dead birds (jittered for clarity) that were ringed in their first year of life (1Y) in the respective study regions (marked by black squares). Dot colours represent the date of death, with blue shades indicating earlier deaths and green-yellow shades indicating later deaths. Three recoveries of dead birds ringed in the northern Netherlands are located outside the mapped extent: one in Ireland and two in the United Kingdom. **c & f**, Annual number of Sandwich terns ringed as 1Y (1995–2021) in the southern Netherlands & Belgium (c) and the northern Netherlands (f). Different grey shades represent different ring types. **d & g**, Region-specific apparent survival estimates of Sandwich terns ringed as 1Y across five age classes in the southern Netherlands & Belgium (d) and the northern Netherlands (g) (see also Supplementary Table **S**3). **e & h**, Estimated number of ringed birds alive per cohort in the southern Netherlands & Belgium (e) and the northern Netherlands (h) at the start of the HPAI outbreak in May 2022. Note that the y-axis scale differs by an order of magnitude with (c) & (f).

Generalised linear mixed models showed that predicted mortality increased significantly with age in both regions. Although both regional and total mortality of ringed birds was consistently higher in the southern Netherlands & Belgium, this geographic trend was not statistically significant (*P*_*reg*_ = 0.056, *P*_*tot*_ = 0.54). Predicted mortality rates ranged from 6.7–10.5% in 4Y birds to 33.2–49.5% in 28Y birds in the southern Netherlands & Belgium, and from 4.9–9.6% to 26.1–47.1% in the northern Netherlands, respectively (Fig. 2). Each additional year of age was associated with an 8.4% increase in the odds of mortality in the regional mortality model (OR = 1.084, 95% CI: [1.056–1.112], *P* = 1.77 × 10^−9^) and a 9.3% increase in the total mortality model (OR = 1.093, 95% CI: [1.068–1.118], *P* = 4.89 × 10^−14^). Both models likely underestimate absolute mortality, since many birds that died, both within and outside the study regions, were probably not recovered or reported. However, the estimated age effect should remain robust, as the probability of recovery is unlikely to show a systematic age-related bias.

**Fig. 2.**
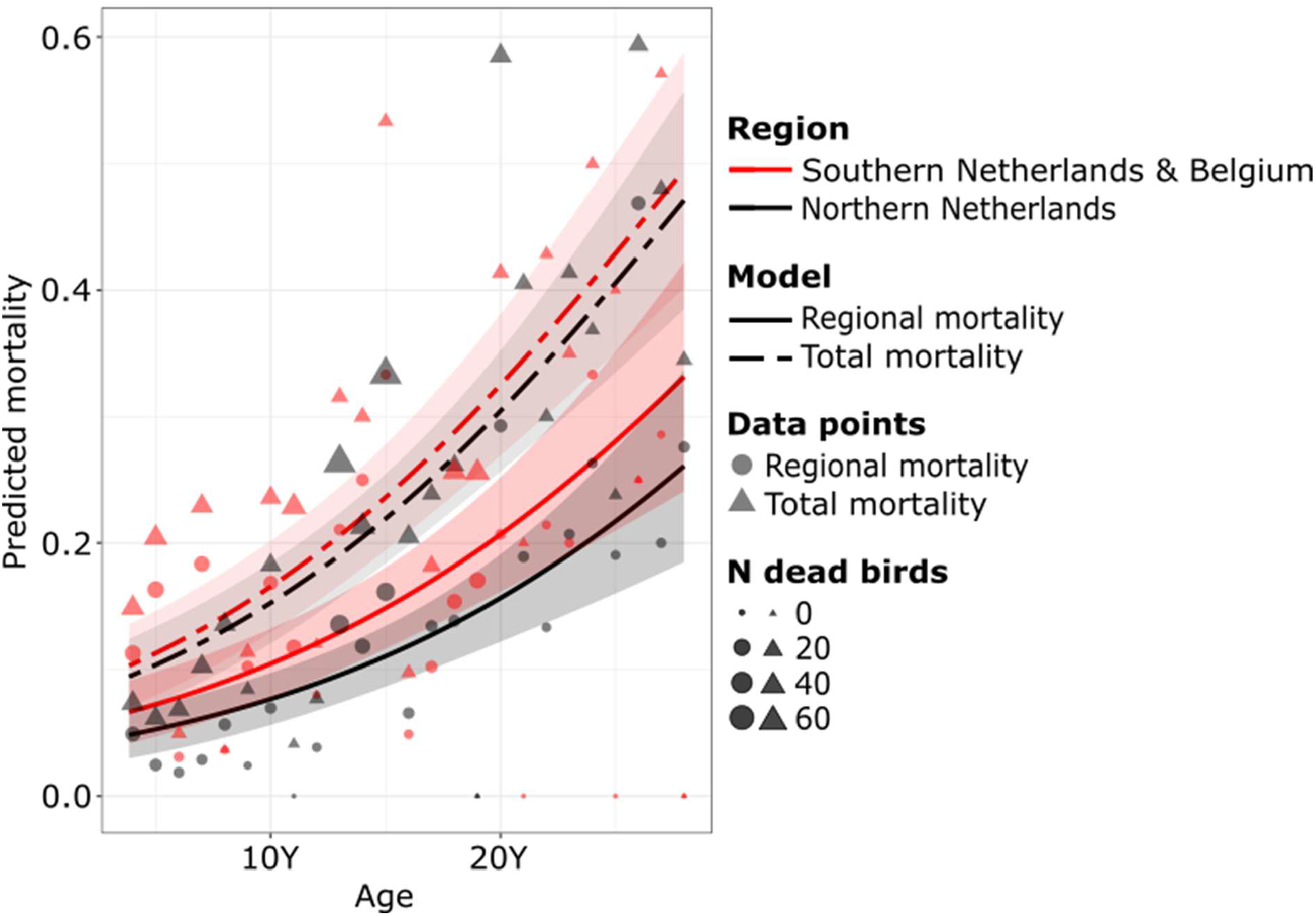
Predicted HPAI-related mortality of Sandwich terns during the 2022 breeding season (May–August) as a function of age and natal region. Data points represent the proportion of birds found dead in 2022 relative to the number of alive birds at the beginning of the HPAI-outbreak. Symbol size scales with the number of dead individuals per year of age. Two models were fitted: (i) a ‘regional mortality model’ including only birds recovered dead within their natal region (solid lines, dots), and (ii) a ‘total mortality model’ including all recoveries of Belgian and Dutch ringed birds in Europe, both in and outside their natal region (dashed lines, triangles). Predicted effects and 95% confidence intervals were estimated using a generalised linear mixed model (GLMM) with a beta-binomial error structure to account for overdispersion. Results are shown separately for the southern Netherlands & Belgium (black lines and symbols) and the northern Netherlands (red lines and symbols). Birds up to their 3Y were excluded from the analysis, as most remain in the wintering areas or arrive late in the colonies. Birds over 28Y were also excluded because many were ringed with aluminium rather than steel rings, which are more likely to be lost and could therefore confound the analysis.

Birds found outside the study regions died significantly later in the season than those found inside (Figs. 1a and 1b; median: 27 June outside, 15 June inside; Wilcoxon rank-sum test, W = 33,991, *P* < 2.2 × 10^−16^; see also^15^). While this temporal pattern may partly reflect a delayed start of carcass removal in some colonies, it probably also reflects that birds relocated to colonies outside the study region after losing their mate or witnessing colony collapse, and subsequently died during or after these movements. Ring resightings indeed confirm that many individuals relocated to foreign colonies, with some initiating a second breeding attempt^17^. As these birds were initially present in the study region but died elsewhere, relocation provides an additional source of downward bias in regional mortality estimates.

Age- and sex-related mortality of seabirds has been documented in relation to fisheries bycatch^18^, migration^19^, wind farm collisions^20^, oil spills^21^, attraction to artificial lights^22^, food stress^23^ and environmental conditions^24^. These patterns are generally attributed to differences in dispersal strategies, at-sea distribution, foraging behaviour, or morphology. Mass mortality caused by HPAI in breeding seabirds is a recent phenomenon, and selective mortality in these outbreaks may have arisen from differences in behaviour or physiology.

To explore whether age differences in phenology contributed to differential mortality, we analysed arrival times of ringed birds in the northern Netherlands, one of the hardest hit regions in 2022 and with well-monitored colonies. First-time breeders (4–5Y) typically arrived later than older birds (linear model, *β* = 23.63 ± 1.97, *t* = 11.98, *P* < 0.001), but still well before the outbreak began (Extended Data Fig. 1). For older birds (6–10Y), there was no relationship between age and date of first sighting (linear model, *β* = 0.43 ± 0.88, *t* = 0.49, *P* = 0.63). Their median arrival date consistently fell between 15 and 18 April, characteristic of the species’ breeding behaviour and more than a month before the first detected cases of HPAI virus infections. This suggests that phenology likely played a minor role in the observed differential mortality. An alternative explanation is immunosenescence, the gradual decline in immune function with age, although research in wild birds remains limited^25^. In vivo studies of HPAI H5 viruses in domestic birds and waterbirds suggest reduced mortality with age^26^, and no age-related immune decline was found in the closely related common tern (*Sterna hirundo*)^27^. These findings contrast with our results and underscore the need for further research on immunosenescence and its potential role in shaping responses of wild birds to HPAI virus infections.

Our findings that HPAI disproportionately affected older, more experienced, and potentially more productive individuals in a long-lived seabird species are deeply concerning, but the long-term consequences for population recovery remain difficult to predict. For example, while instantaneous mortality during outbreaks in the Belgian and Dutch colonies was estimated at 26.7% based on local dead bird recoveries – with true mortality likely to be higher^28^ – the observed decline in numbers of breeders in 2023 was less, at 21.4%. Recruitment of non-breeding adults and first-time breeders, density-dependent effects, and migration between regions may mask the short-term demographic effects of HPAI-related mortality^10,29^. This highlights the importance of continued monitoring, not only across the northwestern European population of Sandwich terns, but also at the colony level, to detect delayed demographic effects and shifts in age structure or productivity.

These uncertainties do not negate the significant threat that HPAI poses to seabird populations worldwide, many of which have experienced severe outbreaks resulting in mass mortality since 2022. Seabirds – including penguins, albatrosses, petrels, gannets, gulls, terns and auks – share life-history traits such as delayed maturity, low annual reproductive output, and colonial breeding, making their populations particularly vulnerable, as adult mass mortality events caused by infectious diseases can have lasting demographic consequences. The disproportionate loss of experienced individuals could undermine the reproductive output of affected colonies, delaying recovery and potentially leading to long-term population declines persisting well beyond the outbreak itself. Quantifying the full demographic impact of such selective mortality requires detailed information on age-specific survival and reproduction, and robust demographic modeling to evaluate their consequences for population trajectories and recovery potential. Such demographic shifts may also erode genetic diversity and disrupt colony structure, reducing the species’ ability to adapt to future environmental pressures. For this group of birds, which is already highly threatened by climate change, overfishing, predation, and habitat loss^30^, the emergence of HPAI adds yet another layer of vulnerability to seabirds in an increasingly unstable world.

## Methods

### Collation of ringing data

We compiled metal ringing data of Sandwich terns covering the period between 1 May 1995 and 31 August 2022 from the national ringing schemes of Belgium and the Netherlands. Belgium has only one colony just across the Dutch border. As this colony was abandoned in 2008 and a lot of exchange between Belgium and the southern Netherlands occurs, the data were grouped. Both countries were at the epicentre of the HPAI outbreak in 2022. The starting year was set at 1995, as from that year onward, all Belgian and Dutch birds were ringed with durable steel rings instead of the weaker aluminium ones, which are likely to be lost over time. In addition, we included data from a colour-ringing program that started in 2012.

All data from the different sources were cross-checked for inconsistencies, such as birds being reported alive in one database but dead in another, conflicting dates and locations, or cases where birds were ‘resighted’ after being found dead. Incorrect records were corrected where possible and else removed. Ringing records from outside the focal period (May – July) or outside the study regions were also omitted, as were resightings from outside the study regions, because they conflict with the assumptions of the capture-mark-recapture models. Only birds ringed in their first year of life (indicated as 1Y, mostly birds ringed just before fledging in their natal colony) were retained as only for these birds exact age determination is possible. To minimize the effect of pre-fledging mortality on the survival estimates, we excluded chicks that died before the end of July in the year of ringing, but retained all other dead recoveries for the mortality predictions. Three types of rings were used during the study period: two types of scientific metal rings (standard rings (M) and larger, field-readable or read metal rings (RM)) and colour-rings (CR). In total, the final dataset contained more than 93,000 records: ringing data for 50,646 Sandwich terns ringed as 1Y (40,428 with M, 3,136 with RM and 7,082 with CR), 42,151 unique (excluding multiple observations of the same bird on the same day) live resightings (4,888 of M, 1,027 of RM and 36,236 of CR), and 975 dead recoveries.

### Survival estimates

Mean annual apparent survival rates (*φ, Phi*) before the mass mortality event in 2022 (1995–2021) were estimated for the northern and the southern Netherlands (including Belgium) using Cormack-Jolly-Seber (CJS) models for encounter data. We used the *RMark* package (version 3.0.0) ^31^ in *R* (version 4.5.1) ^32^, which serves as an interface for the *MARK* program^33^. Our goal was to generate detailed age-specific survival estimates that reliably reflect survival trends across ages, while avoiding models that result in inestimable parameters for certain age classes.

To satisfy the assumptions of the CJS model, resightings were restricted to those within the natal region during a fixed recapture period (the breeding season, 1 May – 31 July). Multiple resightings within the same year were pooled.

We used two approaches to evaluate the goodness-of-fit (GOF) of the mark-recapture data. First, we estimated the overdispersion coefficient (*ĉ*) using the parametric bootstrap GOF procedure in *MARK*, by dividing the observed deviance of the global model by the mean deviance from 100 bootstrap simulations. Second, we assessed the fit of the encounter histories to the CJS model using U-CARE^34^ via the *R2Ucare* package (version 1.0.2)^35^.

As is common with large and heterogeneous capture-mark-recapture datasets, most data subsets produced highly significant GOF test results, indicating violations of key CJS assumptions (Supplementary Table S1). These violations are partly due to the species’ ecology: after fledging, immature Sandwich terns remain at the wintering grounds and are essentially undetectable in the breeding areas, leading to inflated *χ*^*2*^ values in the transience test (test3.sr). To account for this and to avoid overfitting, we defined a priori age-structured models with five age classes^36^. These reflect key life history stages: immatures (1–3Y), first time breeders (4–5Y), young adults (6–10Y), medium-aged adults (11–15Y), and old adults (15+Y).

We developed a series of models in which resighting probability was allowed to vary by age class, ring type (reflecting differences in ring readability, with resighting probability for M < RM < CR), and region, with and without interactions with time. Time was grouped into five-year intervals to reduce the risk of overfitting. Detection probability was also influenced by whether a bird was actively breeding in a given year and whether it nested in a colony (and at a nest site) where it could be observed. These factors likely contributed to the high *χ*^*2*^ values observed in the trap-dependence test (test2.ct), indicating that resighting probabilities were not fully independent. To address this, we included a set of models that explicitly accounted for trap dependence^37^. For each of the model structures described above, we fitted a Pledger mixture model to account for unobserved individual heterogeneity in detection probability, potentially due to differences in sex, condition, or nest site^38^.

Overdispersion was accounted for in the calculation of standard errors and in model selection using Akaike’s Information Criterion adjusted for overdispersion (QAICc). Models with a ΔQAICc < 2 were considered equally supported (Supplementary Table S2).

### Age-composition of the ringed population

We multiplied the region- and age-specific apparent survival rates with the annual number of birds ringed (N_ringed_) in each region to calculate how many birds from each cohort (i.e., those born in the same year) were ‘estimated to be alive and present in the region’ in May 2022 (N_alive_). Specifically, we multiplied the annual survival estimates over the previous years for each cohort. For example, N_alive_ for birds in their 9^th^ year of life (9Y) is calculated as:

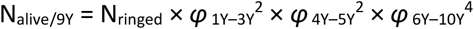

and N_alive_ for birds in their 17Y as:

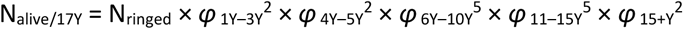

This provided the age composition of the ringed population at the onset of the HPAI outbreak in May 2022.

### Age- and sex-composition of HPAI mortalities

Infection with a HPAI virus often causes acute illness and inability to fly, leading to the discovery of numerous carcasses in or near breeding colonies^28^. In total, 832 birds in their 2Y or older, that were ringed as 1Y in Belgium or the Netherlands (allowing precise age determination), were recovered dead during the 2022 breeding season (May–August) and were presumed to have died from avian influenza. Of these, 595 were found within these two study regions.

Of these, 243 carcasses were collected and sexed between 29 May and 5 July 2022, 82 in the southern and 161 in the northern Netherlands.

### Arrival dates in the northern Netherlands

The dates of first sightings (15 March to 15 July) of 419 Dutch colour-ringed Sandwich terns in their 3– 10Y in the northern Netherlands were used to calculate the median arrival date per cohort. The colonies in the northern Netherlands were surveyed almost daily from mid-March in 2022.

### Statistical analyses

All statistical analyses were performed using *R* (v4.5.1) ^31^. Birds up to their 3Y were excluded from the mortality analysis as 2Y old birds mostly remain in the wintering areas and the median arrival date of birds in their 3Y was 12 days later than the onset of the HPAI epidemic (see Extended Data Fig. 1) and intrinsically had a much lower chance to die in the breeding areas. Birds older than 28Y were excluded because many were ringed with aluminium rather than steel rings, which are subject to rapid wear and subsequent ring loss and could therefore confound the analysis. For the 2022 HPAI outbreak, we modelled the number of dead birds in each cohort, together with the number estimated to be alive and present in the southern Netherlands & Belgium and the northern Netherlands at the start of the outbreak. These two counts were combined using the cbind() function and analysed as the dependent variable in a generalised linear mixed model (GLMM) with a beta-binomial error distribution and a logit link to account for overdispersion, using the *glmmTMB* package (v1.1.10)^39^. Fixed effects included age (covariate ranging from 4Y to 28Y), region (a two-level factor), with and without their interaction, while an observation-level random intercept was included to account for overdispersion. Model selection was based on AIC and likelihood ratio tests, and model fit was assessed using the *DHARMa* package (v0.4.7)^40^.

To account for uncertainty in the spatial scope of mortality detection, we modelled predicted HPAI-related mortality in two ways. First, we included only dead birds recovered in their natal region where they were originally ringed, mirroring the structure of CJS models, where permanent emigration is not distinguished from mortality. This approach yields a conservative estimate of mortality (‘regional mortality’). Second, we used all dead recoveries of ringed birds regardless of where they were found, providing a more inclusive estimate of mortality (‘total mortality’).

We tested whether the sex ratio of adults found dead deviated from the expected 1:1 using a binomial test. To assess the effect of age on the timing of first resighting, we fitted two linear regression models with the Julian day of first resighting as the dependent variable. In the first model, age was included as a five-level factor (6–10Y), and in the second as a two-level factor contrasting younger (4–5Y) and older (6–10Y) birds. We compared date of death between birds found inside and outside the study regions with a Wilcoxon rank-sum test, as both variables were non-normally distributed. Date values were converted to Julian days prior to analysis.

## Data availability

The data to reproduce the results of this study are available on figshare (https://doi.org/10.6084/m9.figshare.30478049).

## Code availability

The code to reproduce the results of this study is available on figshare (https://doi.org/10.6084/m9.figshare.30488306). We provide code implemented in *R* (v4.5.1).

## Acknowledgements

We are very grateful to all people and organizations who contributed data to this manuscript. Ringing and recovery data of metal rings were kindly provided by Henk van der Jeugd (Vogeltrekstation, Netherlands), Didier Vangeluwe (BeBirds, Belgium) and Dorian Moss (EURING). Ringing and colour-ring resighting data were kindly provided by Allix Brenninkmeijer & Date Lutterop. Special thanks to all the volunteers who have given their time to (colour)ringing and to all the ring readers who passed on their observations. We are very grateful to all people collecting data on HPAI mortality and reporting rings of dead birds, with special thanks to Ronald in ‘t Veld for facilitating collection of HPAI victims in the southern Netherlands.

## Author contributions

W.C. designed the study. W.C. collated, summarised and analysed the data with input from U.K., P.v.H. and M.F.. W.C. wrote the first draft of the manuscript with input from U.K. and prepared the final manuscript with input from all authors. All authors approved the final manuscript.

## Additional information

**Extended data** is included in this manuscript.

**Supplementary information** can be found in the accompanying file.

**Correspondence and requests for materials should be addressed to** Wouter Courtens.

## Extended Data

**Fig. 1.**
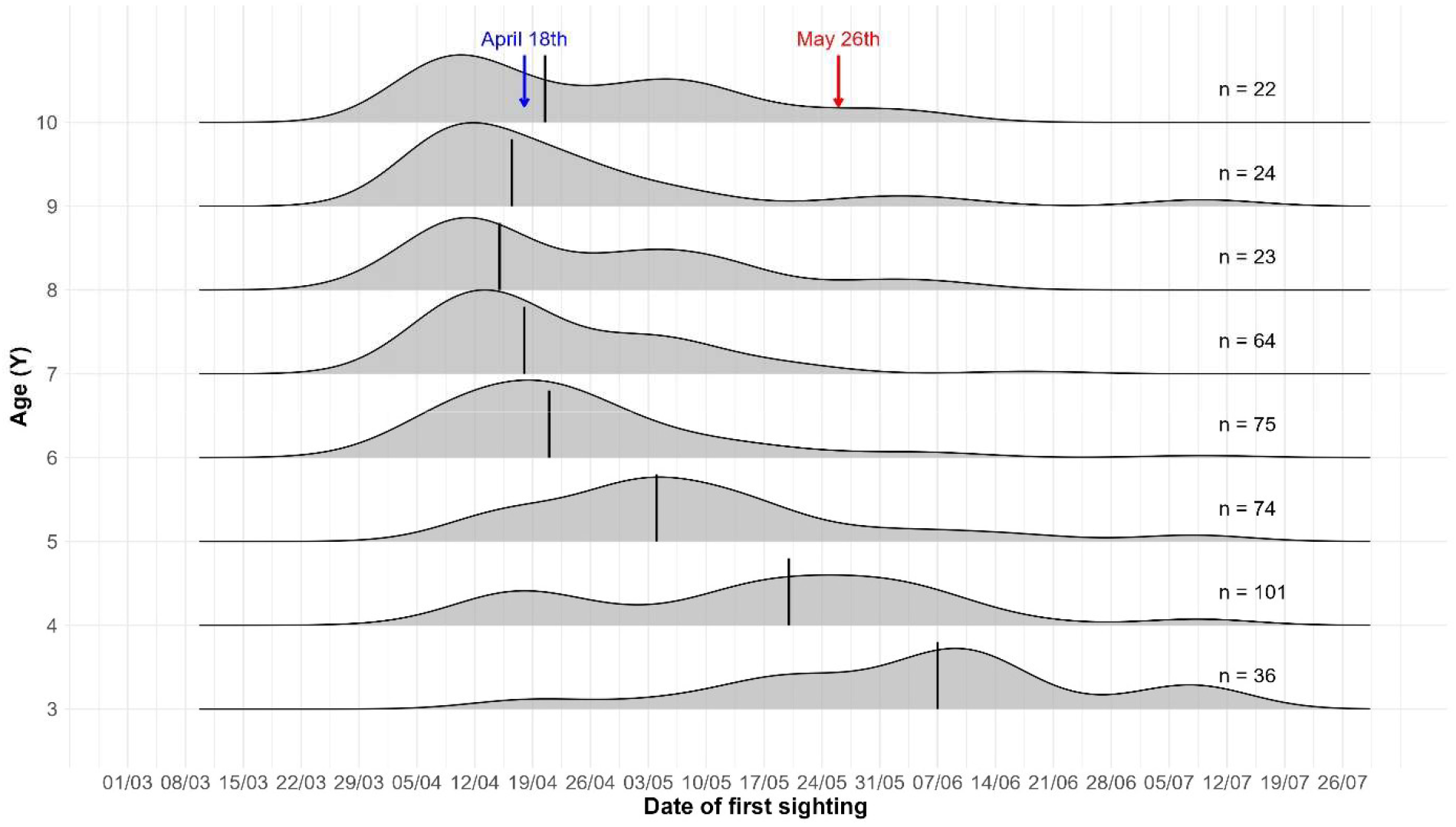
Median date of first sighting of Dutch colour-ringed Sandwich terns of known age in the northern Netherlands. Ridgeline plots show the density of observed birds per date for the age classes 3–10Y. The number of birds observed is shown on the right side of the graph. The blue arrow indicates the median arrival date of birds aged 6–10Y. The red arrow indicates the date when the first HPAI victim was found in the northern Netherlands.

