## Supplementary for "Avian influenza causes age-related mortality in a long-lived seabird"

### Supplementary Information

**Table S1 |** Results of goodness-of-fit tests to examine whether the data fitted a time-dependent Cormack-Jolly-Seber (CJS) model using the *R2Ucare* package implemented in *RMark*. $\chi^2$ is the squared directional statistic, df = degrees of freedom, $\hat{c}$ = overdispersion coefficient (the ratio $\chi^2$/df), *P* = p-value.

| Region | Ringtype | Test | $\chi^2$ | df | *P* | $\hat{c}$ |
|---|---|---|---|---|---|---|
| Southern Netherlands & Belgium | metal | overall | 96.3 | 50 | <0.001 | 1.9 |
| Southern Netherlands & Belgium | metal | test2.ct ('trap dependence') | 26.8 | 12 | 0.008 | 2.2 |
| Southern Netherlands & Belgium | metal | test2.cl | 48.9 | 23 | 0.001 | 2.1 |
| Southern Netherlands & Belgium | metal | test3.sr ('transience') | 20.2 | 12 | 0.064 | 1.7 |
| Southern Netherlands & Belgium | metal | test3.sm | 0.5 | 3 | 0.927 | 0.2 |
| Southern Netherlands & Belgium | colour | overall | 945.5 | 47 | <0.001 | 20.1 |
| Southern Netherlands & Belgium | colour | test2.ct ('trap dependence') | 89.1 | 7 | <0.001 | 12.7 |
| Southern Netherlands & Belgium | colour | test2.cl | 86.6 | 21 | <0.001 | 4.1 |
| Southern Netherlands & Belgium | colour | test3.sr ('transience') | 540.1 | 8 | <0.001 | 67.5 |
| Southern Netherlands & Belgium | colour | test3.sm | 229.6 | 11 | <0.001 | 20.9 |
| Northern Netherlands | metal | overall | 1455.8 | 153 | <0.001 | 9,5 |
| Northern Netherlands | metal | test2.ct ('trap dependence') | 111.9 | 24 | <0.001 | 4.7 |
| Northern Netherlands | metal | test2.cl | 376.5 | 76 | <0.001 | 5.0 |
| Northern Netherlands | metal | test3.sr ('transience') | 829.0 | 24 | <0.001 | 34.5 |
| Northern Netherlands | metal | test3.sm | 138.4 | 29 | <0.001 | 4.8 |
| Northern Netherlands | colour | overall | 1418.0 | 27 | <0.001 | 52.5 |
| Northern Netherlands | colour | test2.ct ('trap dependence') | 101.7 | 5 | <0.001 | 20.3 |
| Northern Netherlands | colour | test2.cl | 99.7 | 10 | <0.001 | 10.0 |
| Northern Netherlands | colour | test3.sr ('transience') | 615.9 | 5 | <0.001 | 123.2 |
| Northern Netherlands | colour | test3.sm | 600.7 | 7 | <0.001 | 85.8 |

**Table S2 |** Model selection showing the 10 best survival ($\varphi$, *Phi*) and recapture (p) models. npar = number of estimated parameters, QAICc = quasi–Akaike Information Criterion corrected for overdispersion, ΔQAICc = difference in QAICc relative to the best-supported model, weight = QAICc weight representing the relative support for each model. AC = age class (1-3Y, 4-5Y, 6-10Y, 11-15Y, 15+Y), reg = region (southern Netherlands & Belgium and northern Netherlands), RT = ring type (metal, read metal and colour), td = trap dependence, mix = mixture model. Time was binned in 5 intervals (1995-2000-2005-2010-2015-2021) to avoid overfitting. The $\hat{c}$-value from the bootstrap GOF test (1.364) was used to adjust standard errors.

| $\varphi$ | p | npar | QAICc | ΔQAICc | weight |
|---|---|---|---|---|---|
| **AC + reg** | **AC + RT + reg * time + td + mix** | 25 | 27972 | 0 | 0.99 |
| AC + reg | AC + RT * time + reg * time + td + mix | 33 | 27981 | 9 | 0.01 |
| AC + reg | AC * time + RT + reg * time + mix | 39 | 27997 | 25 | 0.00 |
| AC + reg | AC + RT * time + reg + td + mix | 29 | 28018 | 46 | 0.00 |
| AC + reg | AC * time + RT + reg + mix | 35 | 28027 | 56 | 0.00 |
| AC + reg | AC + RT + reg * time + mix | 24 | 28084 | 112 | 0.00 |
| AC + reg | AC + RT * time + reg * time + mix | 32 | 28092 | 120 | 0.00 |
| AC + reg | AC + RT * time + td + mix | 28 | 28116 | 144 | 0.00 |
| AC + reg | AC + RT * time + reg + mix | 28 | 28121 | 150 | 0.00 |
| AC + reg | AC * time + RT + mix | 34 | 28325 | 354 | 0.00 |

Across all candidate models, survival ($\varphi$) was modeled as a function of age class and region – $\varphi$(AC + reg). Model selection focused on alternative structures for resighting probability (p). The best-supported model (weight = 0.99) included additive effects of age class and ring type, an interaction between region and time, as well as trap-dependence and a mixture component to account for individual heterogeneity – p(AC + RT + reg * time + td + mix). The second-ranked model, which additionally included an interaction between age class and region in p, received minimal support (weight = 0.01). All remaining models had ΔQAICc > 25 and negligible support. These results show that when survival was consistently modelled for age and regional differences, resighting probabilities were influenced by ring type (reflecting differences in readability of different ring types), temporal and regional interactions (reflecting observer effort changing over time, accessibility of colonies etc.), trap-dependence and individual heterogeneity.

**Table S3 |** Estimates of apparent survival ($\varphi$) from the best model run in *RMark* ($\varphi$ (age class + region) *p* (age class + ring type + region*time + td + mixture)) for five age classes for the northern Netherlands and southern Netherlands with Belgium. $\varphi$ = apparent survival, se = standard error, lcl = lower 95% confidence limit, ucl = upper 95% confidence limit.

| region | age class | $\varphi$ | se | lcl | ucl |
|---|---|---|---|---|---|
| Southern Netherlands & Belgium | 1–3Y | 0.622 | 0.015 | 0.593 | 0.651 |
| Southern Netherlands & Belgium | 4–5Y | 0.985 | 0.019 | 0.844 | 0.999 |
| Southern Netherlands & Belgium | 6–10Y | 0.865 | 0.013 | 0.836 | 0.889 |
| Southern Netherlands & Belgium | 11–15Y | 0.865 | 0.036 | 0.778 | 0.921 |
| Southern Netherlands & Belgium | 15+Y | 0.808 | 0.029 | 0.745 | 0.858 |
| Northern Netherlands | 1–3Y | 0.651 | 0.011 | 0.630 | 0.672 |
| Northern Netherlands | 4–5Y | 0.987 | 0.017 | 0.858 | 0.999 |
| Northern Netherlands | 6–10Y | 0.879 | 0.013 | 0.852 | 0.902 |
| Northern Netherlands | 11–15Y | 0.879 | 0.031 | 0.805 | 0.927 |
| Northern Netherlands | 15+Y | 0.827 | 0.025 | 0.772 | 0.870 |
